# A population genomic unveiling of a new cryptic mosquito taxon within the malaria-transmitting *Anopheles gambiae* complex

**DOI:** 10.1101/2020.05.26.116988

**Authors:** Jacob A. Tennessen, Victoria A. Ingham, Kobié Hyacinthe Toé, Wamdaogo Moussa Guelbéogo, N’Falé Sagnon, Rebecca Kuzma, Hilary Ranson, Daniel E. Neafsey

## Abstract

The *Anopheles gambiae* complex consists of multiple morphologically indistinguishable mosquito species including the most important vectors of the malaria parasite *Plasmodium falciparum* in sub-Saharan Africa. Several lineages have only recently been described as distinct, including the cryptic taxon GOUNDRY in central Burkina Faso. The ecological, immunological, and reproductive differences among these taxa will critically impact population responses to disease control strategies and environmental changes. Here we examine whole-genome sequencing data from a longitudinal study of putative *A. coluzzii* in western Burkina Faso. Surprisingly, many samples are genetically divergent from *A. coluzzii* and all other *Anopheles* species and represent a new taxon, here designated *Anopheles* TENGRELA (AT). Population genetic analysis suggests that GOUNDRY sensu stricto represents an admixed population descended from both *A. coluzzii* and AT. AT harbors low nucleotide diversity except for the 2La inversion polymorphism which is maintained by overdominance. It shows numerous fixed differences with *A. coluzzii* concentrated in several regions reflecting selective sweeps, but the two taxa are identical at standard diagnostic loci used for taxon identification and thus AT may often go unnoticed. We present an amplicon-based genotyping assay for identifying AT which could be usefully applied to numerous existing samples. Misidentified cryptic taxa could seriously confound ongoing studies of *Anopheles* ecology and evolution in western Africa, including phenotypic and genotypic surveys of insecticide resistance. Reproductive barriers between cryptic species may also complicate novel vector control efforts, for example gene drives, and hinder predictions about evolutionary dynamics of *Anopheles* and *Plasmodium*.

## Introduction

Successfully controlling vector-borne diseases will require a comprehensive evolutionary genetic understanding of host species. For malaria, a major global infectious disease afflicting over 200 million people, the relevant vectors are *Anopheles* mosquitoes which transmit *Plasmodium* parasites (Miller, 2002; WHO, 2018). Mosquito-targeting interventions are by far the most effective at reducing malaria infection, substantially exceeding the impact of those that target the parasite directly, like artemisinin combination therapy (Bhatt et al., 2016). However, evolutionary processes such as selection for resistance to insecticides have repeatedly allowed mosquitoes to evade control efforts (Ranson & Lissenden, 2016). Similarly, adaptive resistance is likely to frustrate new control technologies such as gene drives, which involve manipulating mosquito evolution directly (Marshall et al., 2019). Control strategies will need to anticipate and foil these adaptive responses and thus can only succeed if *Anopheles* population genetics is thoroughly understood.

In sub-Saharan Africa, the most important definitive hosts for *Plasmodium* are mosquitoes of the *Anopheles gambiae* species complex. These mosquitoes are among the most genetically diverse animals on Earth (Leffler et al., 2012; *Anopheles gambiae* 1000 Genomes Consortium, 2017). The clade contains nine morphologically-identical species, three of which were only described in the last decade (Coetzee et al., 2013; Barrón et al., 2019). The evolutionary relationships among these cryptic species are complicated due to incomplete lineage sorting and introgression facilitated by porous reproductive barriers (Fontaine et al., 2015). A majority of the genome can cross species boundaries, and this is a frequent and recent phenomenon in response to novel selective pressures such as insecticides (Clarkson et al., 2014; Main et al., 2015; Norris et al., 2015). Additionally, the taxonomic status of some rare yet distinct groups within this complex remains unclear. The subgroup GOUNDRY was identified in Burkina Faso and found to be genetically distinct from its closest known relative, *A. coluzzii* (Riehle et al., 2011; Crawford et al., 2015; Crawford et al., 2016). While GOUNDRY has not been formally described as a separate species, it is genetically distinct from any known species. Further characterization has been impeded because only larval stages have been collected in the field and collecting additional GOUNDRY individuals has proven difficult. There is a substantial need to better understand patterns of gene flow and partitioning of genetic diversity within the *A. gambiae* complex, in order to better predict and mitigate the inevitable evolutionary counterstrategies to vector control efforts.

In this paper, we use whole genome sequencing data to identify yet another new cryptic taxon within the *A. gambiae* complex, occurring in a country (Burkina Faso) where many previous surveys of anopheline mosquitoes have occurred. Though this taxon is closely related to *A. coluzzii*, it shows substantial yet incomplete reproductive incompatibility with it. This new taxon, *Anopheles* TENGRELA (AT), clarifies the origin of GOUNDRY and illuminates the complicated interplay between migration and isolation that characterizes these mosquitoes.

## Materials and Methods

### Samples and sequencing

We chose 287 samples of putative *A. coluzzii* from larval collections in Tengrela, Burkina Faso (10.7° N, 4.8° W) (Table 1). All samples were reared to adults and typed as *A. coluzzii* females based on morphology (Gillies & De Meillon, 1968) and standard molecular assays (Santolamazza et al, 2008). They comprised a longitudinal series across four years (2011, 2012, 2015, and 2016) which were examined as part of a study on insecticide resistance evolution. We extracted DNA from individual mosquitoes using a Qiagen DNeasy Blood & Tissue Kit (Qiagen) following manufacturer’s instructions. We sequenced whole genomic DNA with 151 bp paired-end reads on an Illumina HiSeq X instrument at the Broad Institute, using Nextera low-input sequencing libraries.

**Table 1.** Samples examined.

| Data Source | Year | Sampling | Taxon | N |
| --- | --- | --- | --- | --- |
| This study | 2011 | Tengrela:<br>puddles and/or<br>rice paddies | AT | 51 |
|  |  |  | *A. coluzzii* | 20 |
|  |  |  | Unidentified | 1 |
| This study | 2012-2016 | Tengrela: rice<br>paddies | AT | 0 |
|  |  |  | *A. coluzzii* | 208 |
|  |  |  | Unidentified | 7 |
| Crawford et<br>al., 2016 | 2007-2008 | Central<br>Burkina Faso | GOUNDRY | 12 |
|  |  |  | *A. coluzzii* | 10 |

### Analysis

All reads were aligned to the *Anopheles gambiae* PEST reference genome (assembly AgamP4; Holt et al., 2002; Sharakhova et al., 2007) using bwamem (Li & Durbin, 2009) and variants were called using GATK (McKenna et al., 2010). Initial analysis was restricted to samples with at least 8x median coverage, and AT was identified as distinct from *A. coluzzii* using principal component analysis in R (v. 3.6.2; R Core Team, 2019). Lower-coverage samples were subsequently designated as AT or *A. coluzzii* based on markers that differentiate these taxa in the high-coverage samples; eight samples (coverage 0-2x) could not be unambiguously assigned to taxon and were subsequently ignored, leaving 279 acceptable samples.

Standard statistical tests and data visualization were performed in R (v. 3.6.2; R Core Team 2019). Population genetic statistics such as F_ST_, π, and Dxy were calculated with Perl scripts incorporating the Perlymorphism scripts (Bio::PopGen, BioPerl version 1.007000; Stajich & Hahn, 2005). Statistics were examined in overlapping sliding windows of 1 Mb or 100 kb. Phylogenetic analysis employed RAxML with -m GTRCAT (Stamatakis 2006). *Anopheles* genomes used in phylogenetic analysis (other than AT and *A. coluzzii* from Tengrela) were: *A*. *coluzzii* from elsewhere in Burkina Faso (ERS224009, ERS224023, ERS223804, ERS223963, ERS224782, ERS223946), *A. gambiae* (PEST reference genome, ERS223759, ERS224149, ERS223976, ERS224154, ERS224132, and ERS224151) *A. arabiensis* (SRR3715623 and SRR3715622), *A. quadriannulatus* (SRR1055286 and SRR1508190), *A. bwambae* (SRR1255391 and SRR1255390), *A. melas* (SRR561803 and SRR606147), and *A. merus* (SRR1055284). Annotation and analysis of key genomic regions and alleles was facilitated by VectorBase (Giraldo-Calderón et al., 2015) and the Ag1000G genomic resources (*Anopheles gambiae* 1000 Genomes Consortium, 2017).

In order to directly compare our samples with GOUNDRY, we examined the previously published whole-genome sequences of GOUNDRY and *A. coluzzii* (Crawford et al., 2016; BioProject PRJNA273873). We then examined a representative dataset of 51 AT genomes, 51 *A. coluzzii* genomes from Tengrela, 12 GOUNDRY genomes, and 10 *A. coluzzii* genomes from GOUNDRY habitats (Kodougou and Goundry). To minimize artifacts due to read alignment, we trimmed our Tengrela reads to 100 bp to match the GOUNDRY data, and then aligned all reads to the PEST reference genome and called genotypes jointly as above. For analysis, we removed any variants with missing genotypes, and this jointly-called and filtered dataset was used for a principal component analysis (PCA), ADMIXTURE analysis (Alexander, Novembre, & Lange, 2009), and demographic analysis with *dadi* (Gutenkunsk, Hernandez, Williamson, & Bustamante, 2009). For ADMIXTURE, we chose the value of K with the lowest cross-validation error as recommended. For *dadi*, we also filtered out the 2La inversion and the X chromosome, resulting in a high-quality dataset of 447,181 sites. Heterozygosity in our full dataset is 75x higher than in this filtered dataset, and we estimate that the full dataset represents 85% of the genome, with the rest occurring in long stretches (over 1kb) without called genotypes that may be inaccessible to our genotyping pipeline; thus we adjusted our estimate of nucleotide diversity upwards accordingly when inferring effective population size. We assumed a mutation rate of 3×10^−9^ (Keightley, Ness, Halligan, & Haddrill, 2014), and ten generations per year. We ran multiple models, including models without migration, without population size changes, or with only two periods of differing demographic parameters. Critically for this analysis, we ran models identical to the preferred model but with GOUNDRY originating entirely from either AT or *A. coluzzii*; rejecting these models thus demonstrates that GOUNDRY is admixed.

### Amplicon genotyping

We designed and tested an amplicon-based genotyping method to identify AT in additional samples. We selected 50 diagnostic SNPs and small indels, each with a non-reference allele that is fixed in all AT samples but absent in Tengrela *A. coluzzii* and the Ag1000G data (*Anopheles gambiae* 1000 Genomes Consortium, 2017). For each of these, we designed a primer pair to amplify a PCR product 160 to 230 bp in size using BatchPrimer3 (You et al., 2008; Supp Table 1). We ensured that primers did not overlap common polymorphisms found in *A. coluzzii* or *A. gambiae* (*Anopheles gambiae* 1000 Genomes Consortium, 2017). We ordered primers with the Nextera Transposase Adapters sequences added to the 5′ end in order to eliminate the initial ligation step in library preparation (primer 1: 5′ TCGTCGGCAGCGTCAGATGTGTATAAGAGACAG-[locus specific sequence]; primer 2: 5′ GTCTCGTGGGCTCGGAGATGTGTATAAGAGACAG-[locus specific sequence]).

Primers were tested individually and in various pools of primer pairs to amplify jointly in multiplex PCR using known samples of AT, *A. coluzzii*, and *A. gambiae*. We PCR-amplified 2ng of each DNA sample using the Veriti 96-well Fast Thermal Cycler (Applied Biosystems, Waltham, Massachusetts) with 12.5uL (62.5 U) of Multiplex PCR Master Mix (Qiagen, Hilden, Germany), 2.5 uL of the pre-mixed primer pool (200 nM), and 8 uL of nuclease-free water. The PCR conditions were as follows: 95 °C for 15 minutes; 30 cycles of 94 °C for 30 seconds and 60 °C for 90 seconds, 72 °C for 90 seconds; and 72 °C for 10 minutes. Initial confirmation of correctly sized amplicons was done by gel electrophoresis.

We performed a second round of PCR to attach i5 and i7 Illumina indices (Nextera XT Index Kit v2 set A; Illumina, San Diego, California) to the PCR amplicons. The second round PCR reaction included 5 uL of 1X Platinum SuperFi Library Amplification Master Mix (Thermo Fisher, Waltham, Massachusetts), 5 uL of the first round PCR product, and 1 uL of premixed i5/i7 index primers. The PCR conditions were as follows: 98 °C for 30 seconds; 10 cycles of 98 °C for 15 seconds, 60 °C for 30 seconds, and 72 °C for 30 seconds; and 72 °C for 1 minute. Confirmation of amplification was done by gel electrophoresis.

Amplicon libraries were purified using Agencourt AMPure XP beads (Beckman Coulter, Brea, California) at 1.8x with a final elution volume of 35 uL. Library concentrations were determined using Qubit HS DNA kit (Thermo Fisher, Waltham, Massachusetts). Library concentration and size was confirmed using the Agilent 2100 Bioanalyzer instrument. Libraries were pooled in equimolar concentration and diluted to 180 pmol. A 15% PhiX spike in was added to the diluted pool. We loaded 20uL of the final pool onto an Illumina iSeq 100 instrument at the Broad Institute per the manufacturer’s protocol and sequenced these with 151 bp paired-end reads.

Genotype could be inferred from the resultant reads without an alignment step, simply by counting reads containing diagnostic kmers (24-33bp) overlapping the target polymorphism (Supp Table 1). After determining that a pool of 5 primer pairs sequenced by iSeq was sufficient for taxon identification, we used it to amplify and sequence a novel set of 79 putative *A. coluzzii* samples, alongside a known Tengrela *A. coluzzii* control and 16 empty well negative controls.

## Results

### A new cryptically isolated lineage

In whole-genome sequencing data of 279 putative *A. coluzzii* samples from Tengrela, Burkina Faso (median coverage = 16x; dataset Tennessen et al., 2020), collected across four years, 51 samples were unexpectedly distinct (Figure 1A). The divergent samples, here designated *Anopheles* TENGRELA (AT), were all collected as larvae in 2011, the only year in which sample collection included temporary puddles in addition to rice paddies. AT is not a close match to any known sequenced samples of *A. coluzzii* or *A. gambiae* (*Anopheles gambiae* 1000 Genomes Consortium, 2017). It is similar but not identical to GOUNDRY, which was also described from temporary puddles in Burkina Faso, from sites 250-550 km from Tengrela (Riehle et al., 2011). The genetic divergence from sympatric *A. coluzzii* samples collected simultaneously suggests a strong reproductive barrier.

**Figure 1.**
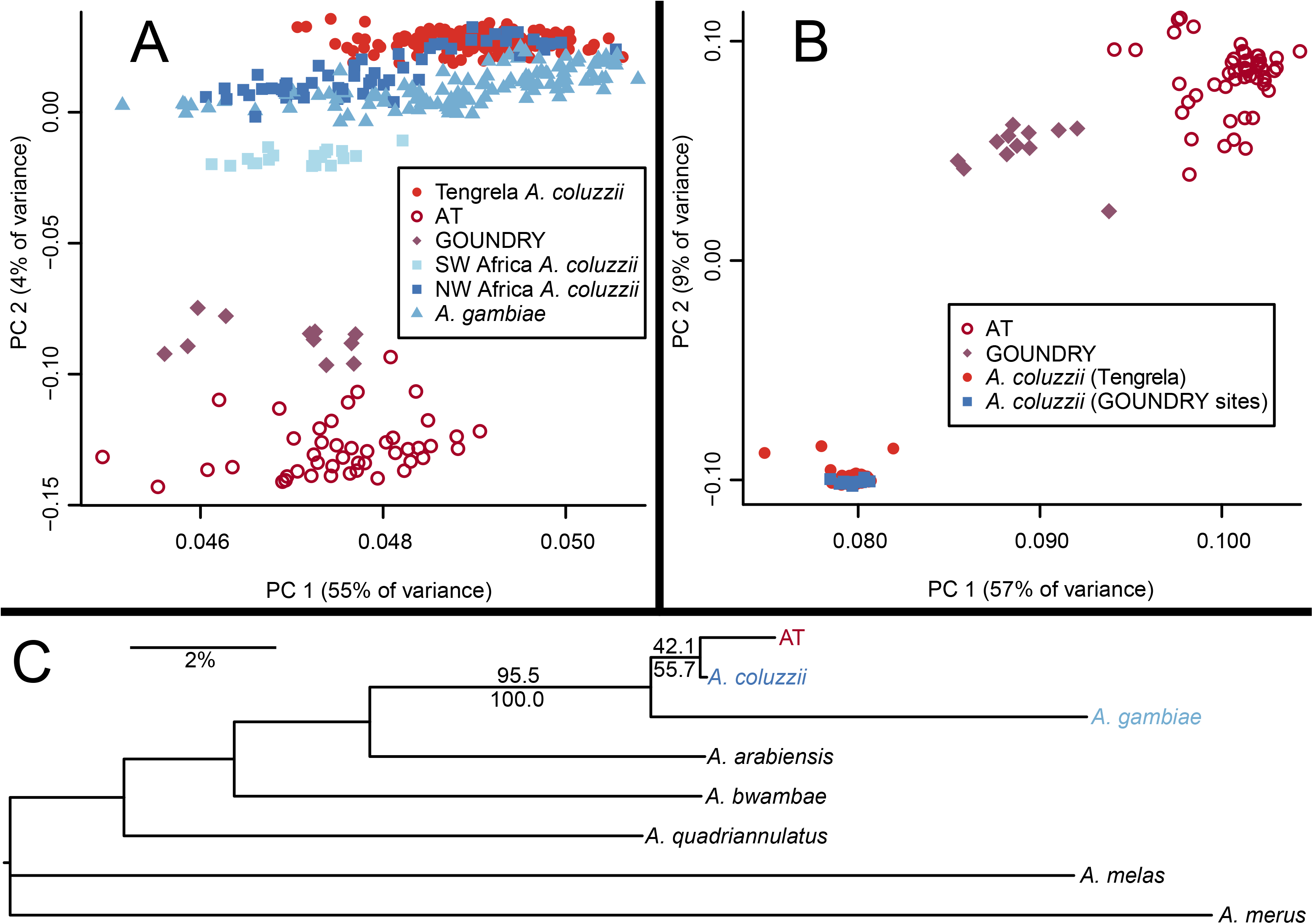
Genetic distinctiveness of AT. (A) In a PCA plot with Tengrela, GOUNDRY, and Ag1000G samples, AT occurs as a distinct cluster close to GOUNDRY. (B) AT remains distinct in a PCA after combining Tengrela samples with GOUNDRY and the *A. coluzzii* samples collected alongside GOUNDRY. To control for differences between studies, all reads were trimmed to the same length, and then alignment and genotyping were performed jointly. (C) Most common phylogenetic topology among sections of AT genome and nominal species of the *A. gambiae* complex. Numbers at branches are not bootstraps, but the percentage of 100 kb windows that support each clade (above branches: entire genome; below branches: X chromosome only). AT is sister to *A. coluzzii* across 42.1 % of the genome (55.7% of the X chromosome), more often than to any other species, and 95.5% of the genome (100.0% of the X chromosome) supports a clade with AT, *A. coluzzii* and *A. gambiae* to the exclusion of the other species.

To further investigate the distinction between AT and GOUNDRY, we jointly analyzed our Tengrela data alongside whole genome sequence data from GOUNDRY and sympatric *A. coluzzii* samples (Crawford et al., 2016). To account for differences in sequencing platforms and genotyping algorithms, we trimmed all reads to 100 bp and jointly aligned them and called genotypes. While *A. coluzzii* from Tengrela and *A. coluzzii* from the GOUNDRY sites were genetically indistinguishable, AT and GOUNDRY remained distinct from each other (Figure 1B). Thus, the distinctiveness of AT is not owing to genotyping artifact, and the divergence between AT and GOUNDRY is greater than that seen for *A. coluzzii* populations over the same physical distance. We therefore infer that AT is not simply an additional population of GOUNDRY.

To examine the relationship between AT and the broader *A. gambiae* species complex, we divided the genome into 100 kb windows and ran a phylogenetic analysis with seven nominal species (Figure 1C). The AT genome is typically sister to *A. coluzzii* (42.1% of windows), *A. gambiae* (33.1% of windows), or to a clade containing only *A. coluzzii* and *A. gambiae* (20.4% of windows), with only 4.5% of windows displaying an alternate topology (Supp Figure 1). The relationship between AT and *A. coluzzii* is especially strong on the X chromosome (Figure 1C; Supp Figure 1), which in *Anopheles* is thought to reflect phylogenetic signal more accurately than the autosomes (Fontaine et al., 2015). As *A. coluzzii* is most often the closest species, and much of the similarity to *A. gambiae* is owing to the 2La inversion (Supp Figure 1), we consider *A. coluzzii* to be the sister taxon to AT for all subsequent analyses.

### GOUNDRY derives from AT and A. coluzzii

Analysis of AT, *A. coluzzii* from Burkina Faso, and GOUNDRY with ADMIXTURE (Alexander et al., 2009) suggests two ancestral populations (K=2; Figure 2A). Populations 1 and 2 are closely approximated by the contemporary AT and *A. coluzzii* populations, respectively (AT: mean population 1 ancestry = 97.1%, range = 71-100%; *A. coluzzii*: mean population 1 ancestry = 0.5%, range = 0-12%). In contrast, GOUNDRY is admixed with approximately equal ancestry in both populations (mean population 1 ancestry = 58.7%, range = 49-74%).

**Figure 2:**
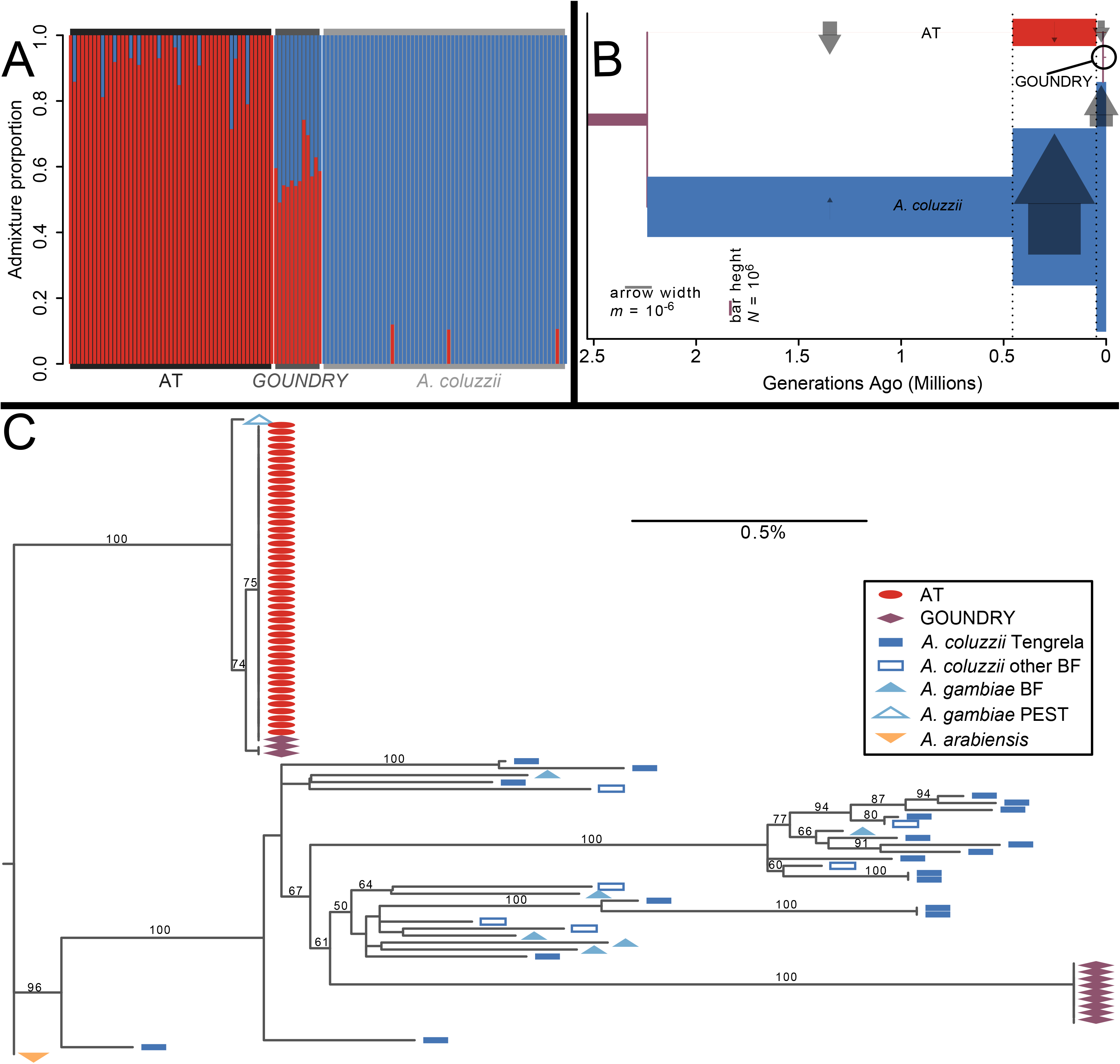
Relationships between AT, GOUNDRY, and *A. coluzzii*. (A) Analysis with ADMIXTURE suggests two ancestral populations, closely approximated by contemporary AT and *A. coluzzii*, with GOUNDRY showing nearly equal ancestry from both. (B) Analysis with *dadi* corroborates this model, with an AT/*A. coluzzii* split over 2 million generations ago, followed by ongoing gene flow and a recent hybrid origin of GOUNDRY (circled). Population sizes (heights of colored bars) and migration rates (widths of arrows) vary across three time periods (demarcated with dotted lines). (C) The mitochondrial DNA phylogeny shows that AT shares a single haplotype that occupies a unique branch close to the *A. gambiae* PEST reference genome. GOUNDRY samples occur near AT or near *A. coluzzii* and *A. gambiae* samples from Tengrela and elsewhere in Burkina Faso (BF), consistent with an admixed origin.

We confirmed the signal of admixture by fitting demographic models to the two-dimensional site frequency spectrum with *dadi* (Gutenkunst et al., 2009). In our best-fitting model, the lineages of *A. coluzzii* and AT diverged over two million generations ago, maintained a small degree of continuous gene flow, and then hybridized less than 20,000 generations ago to form GOUNDRY (Figure 2B; Supp Table 2). Our model included three different time periods among which population sizes and migration rates were allowed to vary. Consistent with the relatively low nucleotide diversity, the effective population size (*N*) of AT was much smaller than *A. coluzzii* across all time periods. It was over 2000-fold smaller in the earliest and longest period, expanded during the middle time period, and contracted again to be approximately 400-fold smaller than *A. coluzzii* today (4.1×10^4^ and 1.7×10^7^, respectively). Migration rates (*m*) ranged from 10^−8^ to 10^−6^, such that each time period the effective number of migrants per generation (4*m* times *N* of the recipient population) was always large in one direction (10 to 20 migrants) and negligible in the other (6×10^−5^ to 3×10^−1^ migrants). Essentially, there has usually been meaningful gene flow from AT into *A. coluzzii*, but gene flow from *A. coluzzii* into AT has been too low to overcome the effects of drift (4*Nm* < 1), except during the middle period 50-450 generations ago, when the AT population size was large and migration from *A. coluzzii* was substantial. GOUNDRY is admixed with 83% ancestry from AT and 17% ancestry from *A. coluzzii*, with *N* slightly larger than AT (8.3 × 10^4^).

We constructed a mitochondrial phylogeny of AT, GOUNDRY, and representative samples of *A. coluzzii*, *A. gambiae*, and *A. arabiensis* (Figure 2C). All AT samples share the same haplotype, which does not occur in any other taxon except for a single GOUNDRY sample. GOUNDRY samples occur in two clades, the one containing AT and another one nested within *A. coluzzii* and *A. gambiae*, consistent with maternal ancestry from both AT and *A. Coluzzii*.

### Genomic characterization of AT

Mosquitoes in the *A. gambiae* complex are typically identified with established molecular markers, including the intergenic region (IGS) of rRNA (Scott, Brogdon, & Collins, 1993; Fanello, Santolamazza, & della Torre, 2002), indels in a SINE200 retrotransposon (Santolamazza et al., 2008), and “divergence island SNPs” showing nearly-fixed differences among taxa (Lee et al., 2013). Analysis of these regions in AT reveals how this taxon easily goes undetected, both in our initial survey of these samples and possibly in other studies (Table 2). At IGS, AT reliably lacks the *HhaI* restriction site found in *A. gambiae* s. s., and instead harbors the AT dinucleotide typical of *A. coluzzii* (Fanello et al., 2002). At retrotransposon S200 X6.1, AT possesses the 230 bp insertion characteristic of *A. coluzzii* (Santolamazza et al., 2008). Interestingly, a SNP within this indel is nearly fixed between AT and *A. coluzzii*, suggesting that amplification of this region could still be diagnostic if it were sequenced and not merely assessed for band size. At divergence island SNPs, AT appears identical to Tengrela *A. coluzzii*, although such SNPs on chromosome 2L are polymorphic in both taxa.

**Table 2.** Key genomic regions characterized in AT, GOUNDRY, and Tengrela *A. coluzzii*.

| Locus | Description | Chromosome | Position | Allele | AT<br>Frequency | GOUNDRY<br>Frequency | Tengrela *Anopheles*<br>*coluzzii* Frequency |
| --- | --- | --- | --- | --- | --- | --- | --- |
| 2La | 22 Mb<br>inversion | 2L | 20000000-42000000 | 2L<sup>+</sup><sup>a</sup> (haplotype) | 48% | 70% | 3% |
| | | | | 2L<sup>a</sup>2L<sup>+</sup><sup>a</sup> (heterozygous<br>genotype) | 73%<br>(expect<br>50%; $\chi^2 =$<br>10.5; $P =$<br>0.001) | 31% (expect<br>42%; $\chi^2 =$<br>36.2; $P <$<br>0.0001) | 5% (expect 5%) |
| Y-linked | Y-chromosome sequence in PEST, excluding sex-determining gene | Y | 50000-230000 | Y sequence present | 94% | 100% | 3% |
| *Rdl* | Insecticide resistance | 2L | 25429235 | SER (resistant) | 32% | 58% | 68% in 2011-2012;<br>38% in 2015-2016 |
| *Kdr* | Insecticide resistance | 2L | 2422652 | PHE (resistant) | 51% | 46% | 78% |
| *CYP9K1* | Insecticide resistance | X | 15242000 | cyp-II (resistant?) | 100% | 67% | 11% |
| *TEP1* | *Plasmodium*<br>resistance | 3L | 11205000 | R (protective) | 0% | 12% | 98% |
| *APL1A* | *Plasmodium*<br>resistance | 2L | 41271000 | APL1A<sup>2</sup> (protective) | 15% | 17% | 80% |
| Xh | 1.5 Mb<br>inversion | X | 8470000–<br>10100000<br>(example<br>SNP at<br>9361641) | Non-reference | 100% | 100% | 0% |
| Sweep<br>region | Unknown<br>phenotypic<br>effect | 3R | 8900000–<br>13300000<br>(example<br>SNP at<br>13081325) | T (non-reference) | 100% | 88% | 0% |
| rDNA IGS | Intergenic region of ribosomal DNA (multiple copies) | UNK (sequence from Scott et al., 1993) | 580-581 | *HhaI* cut site (S-form specific; Fanello et al., 2002) | 0% | 8% | 0% |
| M/S diagnostic SNPs | Divergence island SNPs (Lee et al., 2013) | 2L | 209536, 1274353, 2430786, 2430915, 2431005 | M-form | 46-51% | 54-58% | 22-28% |
|  |  | 3L | 296897, 387877, 413944 | M-form | 99-100% | 100% | 100% |
|  |  | X | 20015634,<br>22105429,<br>22105860,<br>22497157,2<br>2750432,<br>22750572,<br>22944682 | M-form | 99-100% | 96% | 100% |
| S200 X6.1 | 230bp<br>diagnostic<br>indel | X | 22951000 | M-form insertion present | 100% | 100% | 100% |
|  | SNP within<br>insertion | X | 22951586 | T (non-reference) | 100% | 100% | 1% |

Unlike Tengrela *A. coluzzii*, in AT both haplotypes of the 2La inversion are common (Coluzzi et al., 2002; Neafsey et al., 2010), and thus this 22 Mb region represents one of its most striking differences from *A. coluzzii* (Figure 3A). The 2L+^a^ haplotype occurs at a frequency of 48% and shows a remarkable heterozygote excess in violation of Hardy-Weinberg expectations (expected heterozygotes = 25.4, observed heterozygotes = 37; χ^2^ = 10.5; *P* = 0.001), suggesting at least half of homozygous genotypes are selected against. This observation is unexpected, since either the 2L^a^ or 2L+^a^ haplotype can be the major allele across *Anopheles* populations, and thus the inversion is thought to be maintained by geographically-varying selection rather than heterozygote advantage (White et al., 2007). GOUNDRY, in contrast, has a significant heterozygote deficit at 2La (*P* < 0.0001), perhaps owing to inbreeding (Crawford et al., 2016). Our results suggest that the genomic background of AT may facilitate overdominance at this inversion, and thus AT may be an important genetic reservoir which helps to maintain the polymorphism across the genus. Genes potentially under selection in 2La include the highly polymorphic *APL1* gene complex, implicated in immunity against *Plasmodium* (Rottschaefer et al., 2011), and *Rdl*, at which a nonsynonymous Ala-Ser variant conveys resistance to the insecticide dieldrin (Du et sl. 2005). In *APL1*, the protective APL1A^2^ haplotype is rare in AT (15%) but is the major allele in Tengrela *A. coluzzii* (80%). At *Rdl*, resistant Ser occurs at 32% frequency in AT. In Tengrela *A. coluzzii*, this variant represents the largest shift in allele frequency across years (Supp Figure 2), decreasing from 68% in 2011-2012 to 38% in 2015-2016, consistent for selection against costly resistance as organochloride use has declined. While *Rdl* in *A. coluzzii* must occur on the (nearly fixed) 2L^a^ haplotype, in AT it is more often associated with 2L+^a^.

**Figure 3.**
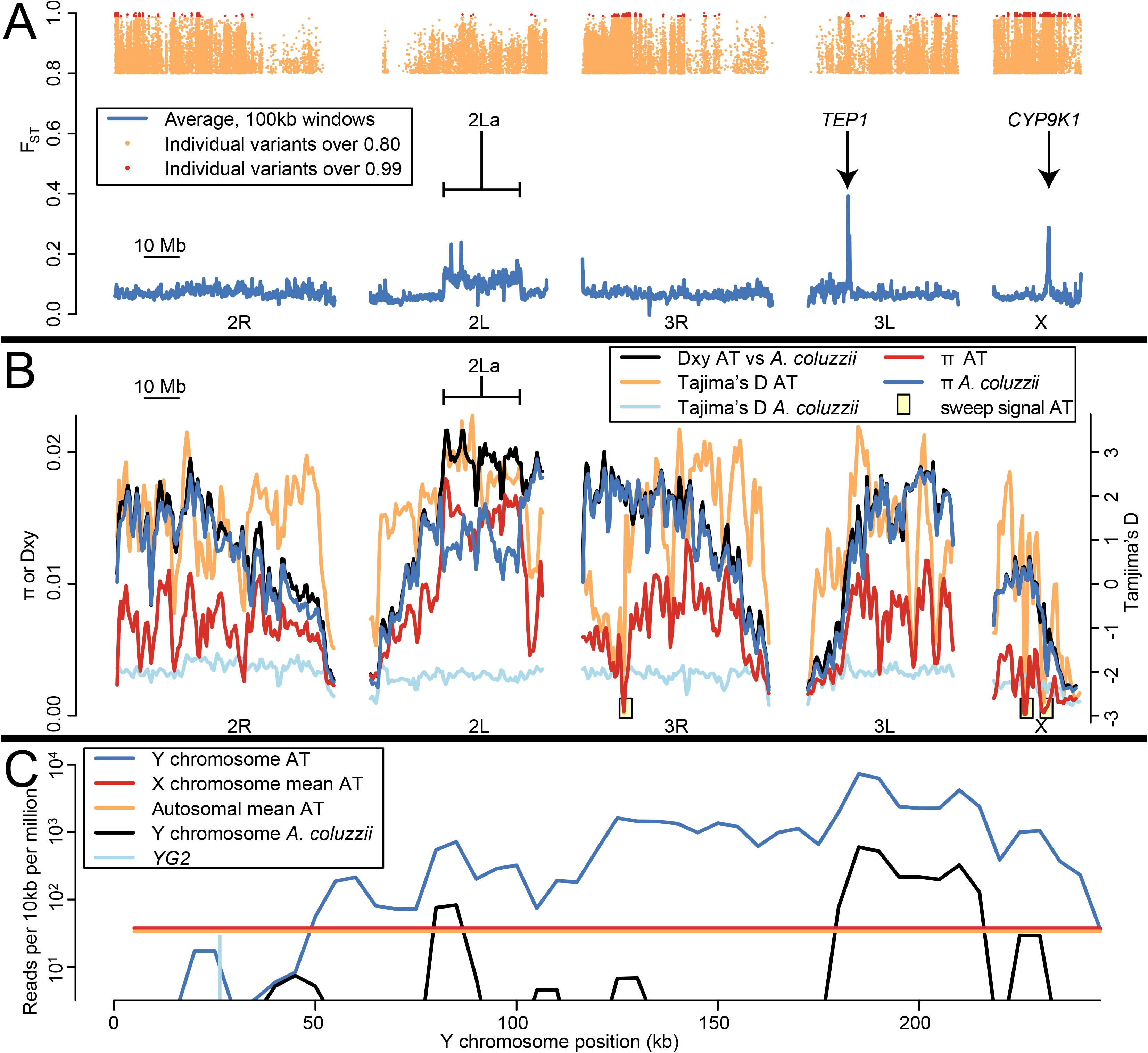
Unique genomic characteristics of AT. (A) F_ST_ across the genome between AT and Tengrela *A. coluzzii*. Tens of thousands of variants distributed across the genome are highly divergent between these taxa (F_ST_ > 0.8; orange dots at top), while over a thousand sites, concentrated in several clusters, are fixed or nearly so (“definitive differences”, F_ST_ > 0.99; red dots at top). Average F_ST_ in 100 kb windows is more modest (blue lines), but three regions stand out representing the 2La inversion, *TEP1*, and *CYP9K1*. (B) Nucleotide diversity (π), intertaxon divergence (Dxy), and site frequency spectra (Tajima’s D) in AT and *A. coluzzii*. Nucleotide diversity is low in AT except at the 2La inversion. Tajima’s D is mostly positive in AT, but three regions show low Tajima’s D, low π, and many high-F_ST_ sites (as shown in A), suggesting selective sweeps. (C) In most AT females, read coverage along most of the reference Y chromosome substantially exceeds the X/autosomal average. Coverage is negligible around sex-determining gene *YG2*. *A. coluzzii* females, in contrast, typically show negligible coverage across the entire Y except for a few repetitive sections.

Several other genomic regions are unusually divergent between AT and *A. coluzzii* (Table 2). Along with 2La, both *TEP1* and *CYP9K1* are outliers with respect to mean F_ST_ between these taxa (Figure 3A). *TEP1* encodes a complement-like immunity protein that occurs in highly dissimilar allelic forms correlated with resistance to *Plasmodium* (White et al., 2011). All *TEP1* alleles in AT are of the S (susceptible) type, while the R (resistant) type is nearly fixed in Tengrela *A. coluzzii*. *CYP9K1* is a P450 gene associated with resistance to insecticide (Main et al., 2015). The cyp-II haplotype of the *CYP9K1* region is fixed in AT and rare in Tengrela *A. coluzzii*; there is suggestive but inconclusive evidence that this allele conveys increased resistance (Main et al., 2015). In contrast to the pronounced divergence at *CYP9K1*, the well-known insecticide-resistance polymorphism *Kdr* (Donnelly et al., 2009) shows similar, intermediate frequencies across both AT and Tengrela *A. coluzzii*.

Nucleotide diversity is substantially lower in AT (π = 0.007) than in *A. coluzzii* (π = 0.012) (Figure 3B). This trend is consistent across the genome except within the 2La inversion (AT: π = 0.015 in 2La, π = 0.006 elsewhere; *A. coluzzii*: π = 0.013 in 2La, π = 0.012 elsewhere). This difference is also consistent among individuals, such that the range of heterozygosity per individual for AT does not overlap that of *A. coluzzii* (Supp Figure 3). Across the genome, divergence between these taxa as measured by Dxy closely matches π in *A. coluzzii*, as this taxon that contributes the majority of variation; the exception occurs in 2La, when both Dxy and AT π are maximined. The site frequency spectra of AT and *A. coluzzii* are also quite distinct. In *A. coluzzii*, Tajima’s D is very negative (D = −2.0) and fairly uniform across the genome (SD = 0.2), reflecting its recent population expansion (Figure 2B; *Anopheles gambiae* 1000 Genomes Consortium, 2017). In contrast, Tajima’s D in AT is positive on average (D = 1.3), consistent with a recent dramatic decrease in population size (Figure 2B), but it’s also quite variable across the genome (SD = 1.3). Three genomic regions in AT, comprising about 2% of the genome, show a combination of highly negative Tajima’s D and unusually low nucleotide diversity, and together they harbor a third of the “definitive differences” (defined here as F_ST_ > 0.99; Figure 3A) between AT and *A. coluzzii*. This suite of signals suggests positive selective sweeps in these three regions in AT (Figure 3B). One putative sweep is observed on 3R. Though the signal extends from approximately 8.9 Mb to 13.3 Mb, it is concentrated in a one-megabase section from 11.5 to 12.5 Mb which contains 125 definitive differences (7% of the genome-wide total) and also the lowest autosomal nucleotide diversity (π = 0.0003) and lowest Tajima’s D genome-wide (D = −2.9). Multiple genes occur in or near this region, with no obvious single selection target. Several of these genes have known phenotypic effects, including *IR21a* which encodes a thermoreceptor implicated in heat-mediated host-seeking (Greppi et al., 2020), and a cluster of cuticular proteins tied to insecticide resistance (Nkya et al., 2014; Huang et al., 2018). The other two selective sweep signals occur on the X chromosome, which even outside of these regions shows lower nucleotide diversity overall than the autosomes. The putative inversion Xh between 8.5 to 10.0 Mb, which also shows a sweep signal in GOUNDRY (Crawford et al., 2016), contains 15% of all definitive differences with *A. coluzzii* and has the lowest nucleotide diversity in the AT genome (π =0.0001). The third signal overlaps *CYP9K1* on the X between 13.5 to 16.0 Mb. It accounts for 10% of all definitive differences and also shows low nucleotide diversity (π = 0.0004). These two sweep regions on X show the lowest Tajima’s D values in the genome outside of the 3R sweep region (Xh vicinity D = −2.6; *CYP9K1* vicinity D = −2.7).

A final major genomic feature of AT concerns the Y chromosome. All AT samples were morphologically typed as female and confirmed as such due to coverage similarity between the X chromosome and autosomes. However, 94% of them showed high coverage across the majority of the Y chromosome, except for the first 50 kb which includes the sex-determining region and sex-determining gene *YG2* (Figure 3C). For most of the Y chromosome after 50 kb, coverage exceeded the autosomal and X-chromosome averages by over an order of magnitude, implying that this sequence occurs repeatedly. The most likely explanation is that multiple copies of most of the Y chromosome have merged with an autosome or the X chromosome in AT, consistent with the highly repetitive and dynamic nature of the *Anopheles* Y (Hall et al., 2016).

### A diagnostic protocol

In order to search for AT among additional samples, we developed a diagnostic protocol based on amplicon sequencing. We first identified 50 diagnostic SNPs and small indels that distinguish AT from Tengrela *A. coluzzii* and the Ag1000G samples (Supp Table 1; Supp Fig 4). We tested several pools of primer pairs and found that a pool of five primer pairs could be jointly amplified and sequenced to sufficient coverage on an Illumina iSeq 100. We genotyped each locus by searching the reads for kmers overlapping the target polymorphism. In a set of 10 AT samples and 10 Tengrela *A. coluzzii* samples, each locus yielded an unambiguous genotype in every sample (median number of read pairs per locus per sample = 7905; mean = 9047; range = 954 to 25,180; Fig 4). Specifically, for all loci at all AT samples, more than 90% of read pairs had the AT allele, and for all loci at all *A. coluzzii* samples, more than 90% of read pairs had the *A. coluzzii* allele. The small numbers of incorrect reads could be due to index hopping among these jointly-sequenced samples (van der Valk, Vezzi, Ormestad, Dalén, & Guschanski, 2019) or low-level contamination. Coverage at the negative control was lower but non-negligible (76 to 1460 read pairs per locus, mean = 411.6), presumably due to similar errors. However, for four out of five negative control loci, both alleles occurred in more than 20% of read pairs. These results suggest that taxonomy can be confidently assigned and false positives avoided if two filters are applied. First, samples should be excluded if they have fewer than 10% of the expected number of reads pairs (i.e. fewer than about 4000 read pairs for a typical iSeq lane with 96 samples). Second, for each sample, a majority of loci should implicate the same taxon with at least 90% of reads. Either one of these filters would exclude our negative control.

**Figure 4.**
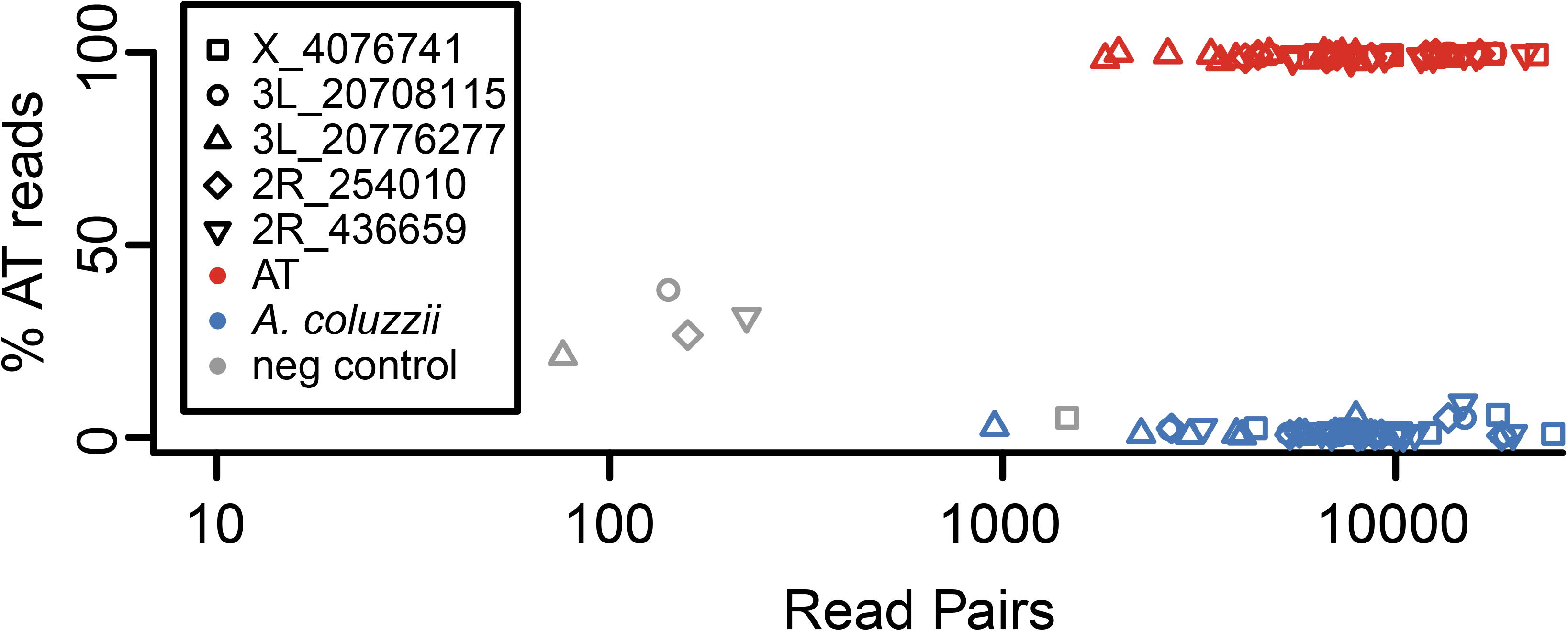
Distinguishing AT from *A. coluzzii* using amplicon genotyping. A pool of five primer pairs were selected that amplify five polymorphisms diagnostic for AT, and jointly amplified PCR products were sequenced (10 AT, 10 *A. coluzzii*, and one negative control). For each marker, the vast majority of read pairs are consistent with the known taxonomic category as inferred from whole genome sequencing, allowing for unambiguous identification. A small number of read pairs were erroneously assigned to the negative control (“neg”), indicating that a conservative test should exclude any sample with unusually low coverage and/or intermediate frequencies of both alleles.

We then used this pool of five primer pairs to amplify and sequence a novel set of 79 putative *A. coluzzii* samples. All negative controls, and three real samples, had fewer than 4000 read pairs each and were discarded by the criteria outlined above. The remaining 76 samples had acceptable coverage (median number of read pairs per locus per sample = 1799; mean = 3466; range = 12 to 25,357) and were all genotyped as unambiguously *A. coluzzii* (maximum observed proportion of read pairs with AT allele: 2.6%). Thus, we failed to detect AT in this novel sample set.

## Discussion

We present a novel taxon within the *Anopheles gambiae* species complex, *Anopheles* TENGRELA (AT). AT is most closely related to *A. coluzzii*, and together these two lineages explain the origin of GOUNDRY via hybridization. AT is genetically distinct from sympatric *A. coluzzii* and shows numerous fixed or nearly-fixed differences across the genome, as well as large difference in genomic architecture such as the 2La inversion polymorphism, the fixed Xh inversion, and the presence of Y-chromosome-associated sequence in females. These differences are presumably maintained by strong reproductive barriers. However, reproductive isolation is not complete, as our demographic analysis indicated ongoing gene flow and a hybridization event within the last few thousand years (Figure 2). Such occasional gene flow is typical among nominal species of the *A. gambiae* complex (Fontaine et al., 2015; Main et al., 2015). Lacking adequate phenotypic and ecological data, we refrain from formally describing AT. However, it represents a unique lineage and is neither nested within *A. coluzzii* nor any other described species, nor is it synonymous with GOUNDRY.

Our demographic model suggests that AT and *A. coluzzii* diverged over two million generations ago, or 200,000 years ago assuming ten generations per year. This result depends on numerous estimates, including mutation rate and the proportion of the genome genotyped accurately, and is therefore imprecise. In general, we endeavored to be conservative in our estimate of differentiation between AT and *A. coluzzii*. For example, we assumed no effect of purifying selection on the variants used in demographic analysis, but if purifying selection has acted it would mean the true time since divergence of AT and *A. coluzzii* is even greater than estimated here. Regardless, our results suggest that cladogenesis predates the rise of agriculture in sub-Saharan Africa and was not driven by adaptation to anthropogenically-disturbed habitats. The population sizes of both lineages increased following this split, bolstered by cross-lineage migration. Then approximately 5,000 years ago AT decreased dramatically in population size while *A. coluzzii* increased further as is well documented (*Anopheles gambiae* 1000 Genomes Consortium, 2017), leading to a modern *A. coluzzii* population hundreds of times larger than AT. If effective population size today approximates census size, the relative rarity of AT could partially explain why it has not been detected previously. Interestingly, the AT population crash occurred around 3000 BCE during the advent of African agriculture (Shaw, 1972), which is hypothesized to have fostered the diversification of the *A. gambiae* complex (Coluzzi et al., 2002; Crawford et al., 2016). Thus, the decline of AT leading to its present-day rarity may have been driven by anthropogenic modifications to habitat, which perhaps favored *A. coluzzii* instead. Our demographic model is similar to the one previously suggested for comparing GOUNDRY and *A. coluzzii* (Crawford et al., 2016). Relative to that model, our estimate of the split with *A. coluzzii* is nearly twice as old, and there are minor differences in population size changes and migration rates. By incorporating AT, we reveal GOUDNRY’s surprisingly recent origin as a distinct lineage.

We know little about the ecology of AT. While GOUNDRY is known from several sites across the hot, arid Sudano-Sahelian region of central Burkina Faso, AT in Tengrela occurs in the cooler, wetter, tropical savannah Sudanese climactic zone of southwestern Burkina Faso. If GOUNDRY is derived from a local AT population, it would suggest that AT possesses a broad tolerance for variable tropical climate. Alternatively, the *A. coluzzii* ancestry of GOUNDRY may have permitted it to colonize more arid climates than AT can tolerate. We only observed AT in 2011, the sole year when samples were collected in temporary puddles and not just rice paddies, which suggests an ecological specificity. As *A. coluzzii* is known to prefer rice paddies in this region of Burkina Faso (Gimonneau et al., 2012), puddle habitats are presumably enriched for AT. Although we cannot rule out immediate local decline or extinction of AT between 2011 and 2012, such a dramatic change seems implausible. Puddle-specificity of AT is also consistent with the known habitat of GOUNDRY (Riehle et al., 2011). The putative selective sweep regions in AT may contain genes that convey unique adaptive features to AT, but these remain to be characterized. Insecticide resistance alleles are present in AT (Table 2), including at *CYP9K1* which occurs within a putative selective sweep region. Selection pressure from regular exposure to insecticide would imply that AT may commonly occur in human-dominated habitats like the Tengrela village. We do not know if adults are anthropophagous, or if they carry *Plasmodium*. The majority of malaria transmission in Africa is due to *A. gambiae* and *A. coluzzii* (*Anopheles gambiae* 1000 Genomes Consortium, 2017), and their close phylogenetic relationship with AT (Figure 1C) suggests similar vectorial capacity. Known *Plasmodium*-resistance alleles tend to be rare in AT (Table 2), though the selection pressures on these immune loci are multifaceted and probably mediated by multiple infectious agents. The substantial vectorial capacity of GOUNDRY (Riehle et al., 2011) is plausibly shared with AT, but live adult AT will be required to pursue this hypothesis further.

Regardless of any direct vectorial capacity, cryptic *Anopheles* taxa have the potential to stymie malaria control efforts in at least three ways. First, reproductive barriers can thwart efforts to eliminate or modify the *A. gambiae* complex via gene drive (Marshall et al., 2019). A gene drive targeting *A. coluzzii* will not necessarily spread to sympatric AT, which might even expand its population size in response to a population crash of *A. coluzzii*. Second, because reproductive barriers are porous, adaptive genetic diversity maintained within AT can be shared with congeners via introgression, enhancing their capacity to survive and evolve. Rare, semi-isolated taxa like AT can thus serve as allelic reservoirs, facilitating adaptation to insecticides, gene drives, or even climate change. For example, the 2La inversion is associated with adaptation to climate (Cheng et al., 2012), and it shows heterozygote advantage in AT. Overdominance in AT could explain the persistence of this polymorphism in a rare taxon with otherwise low nucleotide diversity. In contrast, selection tends to eliminate one other the other 2La haplotype in *A. coluzzii* and *A. gambiae* populations, so AT could be an important genetic reserve for these species if a previously disfavored and purged allele becomes once again beneficial. Finally, misidentified cryptic taxa can confound scientific studies and lead to incorrect inferences about *Anopheles* biology and inaccurate predictions about disease epidemiology and the outcomes of vector control. For example, the *A. gambiae* species complex first came to light following confusing discordances in insecticide resistance phenotype between field and captive mosquito populations (Davidson 1956). Captive mating experiments subsequently demonstrated the discordance was due to the inability to discern *A. gambiae* from another morphologically identical member of the complex, *A. arabiensis* (Davidson 1956, Davidson & Jackson, 1962). More recently, Gildenhard et al. (2019) noted a striking difference in *TEP1* allele frequencies between two populations of *A. coluzzii* in Burkina Faso, which was attributed to ecologically-varying selection. While this is a very plausible and potentially accurate explanation, the results could also be explained by misidentified AT within the samples, as AT has a very different *TEP1* profile from *A. coluzzii* (Figure 3A, Table 2). Since AT and *A. coluzzii* appear similar at standard diagnostic loci (Table 2), this is just one of many studies on *A. coluzzii* that could be potentially confounded by AT.

It will be crucial to correctly identify *Anopheles* taxa in order to draw accurate conclusions that can inform disease control policy. Our amplicon-based diagnostic protocol (Figure 4; Supp Table 1) provides a clear methodology for identifying AT. The pool of five primer pairs can be amplified and sequenced jointly, yielding high accuracy. For further genotyping as needed, we provide primer pair sequences for a total of 50 diagnostic polymorphisms (Supp Table 1; Supp Figure 4), though not all primers have been empirically validated. We encourage *Anopheles* biologists to genotype existing DNA samples, especially of putative *A. coluzzii* from Burkina Faso and adjacent countries, and to seek AT in future surveys. AT will probably not be the last cryptic taxon discovered within this extraordinarily diverse clade. Mapping these intertwined lineages across Africa will be an essential ongoing task with an inestimable impact on human health.

## Acknowledgements

Our thanks to the residents of Tengrela where this study was conducted, and especially volunteer field workers from this village. We are grateful to the field team from CNRFP for helping with mosquito collection. We also thank Sanjay Nagi, Patricia Pignatelli, Natalie Lissenden, and Tim Farrell for assistance with sample collection, molecular preparation, and/or bioinformatics. Data interpretation was aided by helpful commentary from Angela Early, Stephen Schaffner, Aimee Taylor, Seth Redmond, and others. Mosquito collections in Burkina Faso were supported by EC FP7 Project grant no: 265660 “AvecNet” and Wellcome Trust Collaborative Award (200222/Z/15/Z). This project has been funded in whole or in part with Federal funds from the National Institute of Allergy and Infectious Diseases, National Institutes of Health, Department of Health and Human Services, under Grant Number U19AI110818 to the Broad Institute.

## Data Accessibility

Raw Illumina reads from whole-genome sequencing have been deposited in NCBI SRA at https://www.ncbi.nlm.nih.gov/sra.

## Author contributions

JAT performed all data analyses and wrote the paper with the assistance of all co-authors. RK headed the development and testing of the amplicon panel. VAI, KHT, WMG, NS, and HR provided samples and assisted with data interpretation. DEN oversaw the project and assisted with data interpretation.

