## Supplementary Material for "A population genomic unveiling of a new cryptic mosquito taxon within the malaria-transmitting *Anopheles gambiae* complex"

**Tennessen et al. 2020**
**Supplementary Material**

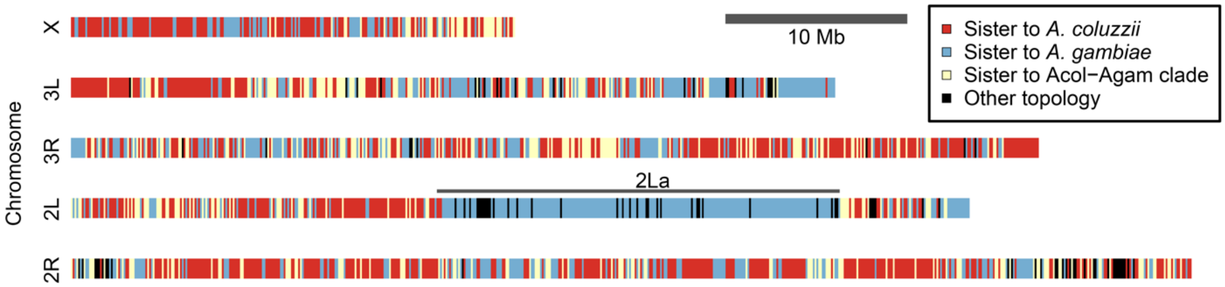


**Supplementary Figure 1. Phylogenetic relationships between 100kb sections of AT genome and nominal species *A. coluzzii*, *A. gambiae*, *A. arabiensis*, *A. merus*, *A. melas*, *A. quadriannulatus*, and *A. bwambae* (Figure 1C). AT is most often sister to *A coluzzii*, the main exception being the 2La inversion (indicated), and AT nearly always forms a clade with *A. coluzzii* and/or *A. gambiae* to the exclusion of the other species.**

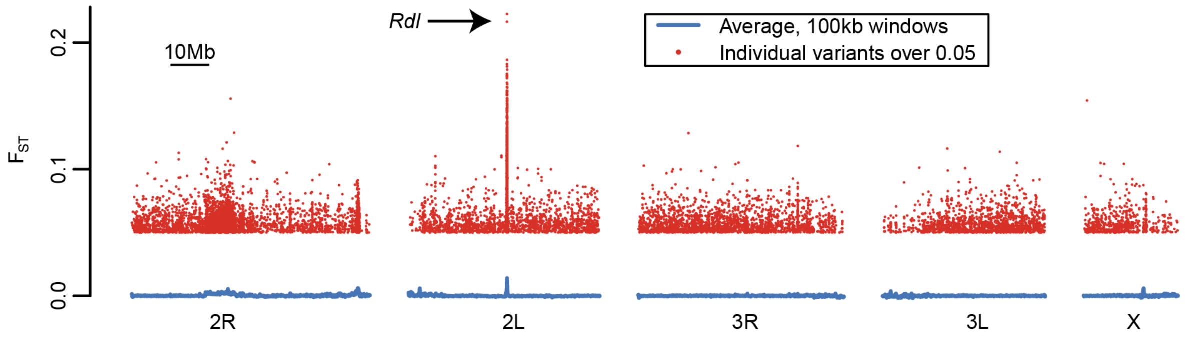


**Supplementary Figure 2. $F_{ST}$ across the genome in Tengrela *A. coluzzii* between early years (2011 and 2012) and later years (2015 and 2016). Average $F_{ST}$ in 100 kb windows is nearly zero between years (blue lines), and only a minority of variants exceed $F_{ST}$ of 0.05 (red dots). The most extreme $F_{ST}$ values, some exceeding 0.2, occur in the vicinity of the *Rdl* gene which affects insecticide resistance.**

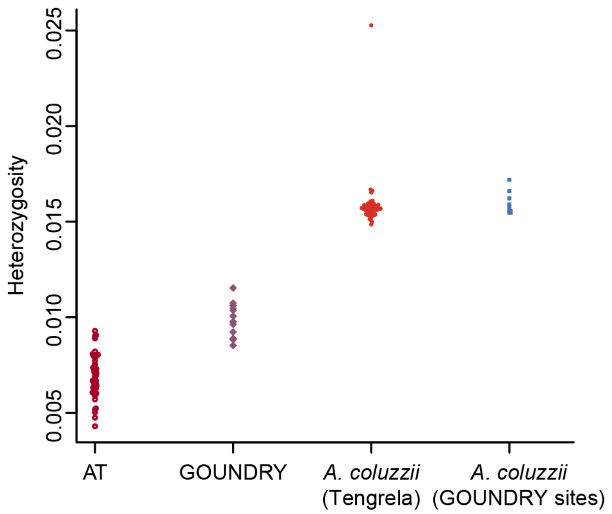


**Supplementary Figure 3. Heterozygosity per individual for the jointly called samples (Fig 1B), excluding 2La. AT mosquitoes are consistently less heterozygous than *A. coluzzii* mosquitoes, with GOUNDRY showing intermediate heterozygosity. The exceptionally high heterozygosity of a single *A. coluzzii* individual (0.025) may be due to contamination, sequencing error, or introgression from an unknown source.**

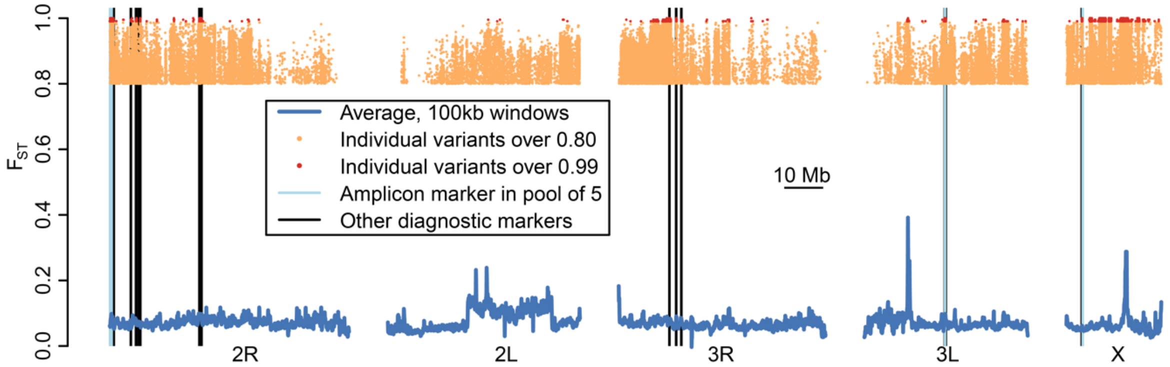


**Supplementary Figure 4. $F_{ST}$ across the genome between AT and Tengrela *A. coluzzii* as in Figure 3A, with diagnostic markers (Supp Table 1) indicated. The 50 diagnostic polymorphisms for which we designed primer pairs are shown, but many are too closely adjacent to be individually discernable at this scale. The five markers used in amplicon genotyping (Figure 4) are indicated in light blue; all are more than 60 kb apart from each other, but the pair on 2R and the pair on 3L cannot be easily distinguished at this scale.**

Supplementary Table 1. Primer pairs targeting diagnostic polymorphisms.

| Polymorphism | Forward | Reverse | Tm °C | Size bp | Pool | AT kmer | *A. gambiae*/*A. coluzzii* kmer |
|---|---|---|---|---|---|---|---|
| X_3930767 | TTCATCATCACTTTTTCTTTTCCTC | TTGAAACGATATGGGCAACA | 60 | 222 | no | na | na |
| X_4076741 | GTGAACCAGAGGTTGCCACT | TTCGCGAGTCTTGAGAGGTT | 60.1 | 228 | yes | AGTNTACCCCANCACAGACGCTTT | AATNTACCCCANCACAGACGCTTT |
| Mt_13083 | AATTTTAATACCACCCCAGTAAAA | TTGAAGGCTGGTATGAATGG | 57.9 | 201 | no | na | na |
| 3L_20669422 | AGCTAATCCGGGGGAGTTTA | CACTGGCCGATTGAGGTTAT | 59.9 | 218 | no | na | na |
| 3L_20708115 | TTTGAGAACGAAGACGAGCA | GAAACATTCCAACCCAGAGC | 59.6 | 228 | yes | TTTTTATCATATCTTACAAATCACAGCAT | TTTTTATCATATCATACAAATCACAGCAT |
| 3L_20770338 | GTCGAACAGTCCGAGCAGAT | TCGGGAAGAACCTCAAAGTG | 60.3 | 230 | no | na | na |
| 3L_20772890 | TGTTGGGAGTGTTGCAGCTA | TGTCCCTCCTAACCCAGATG | 60.2 | 219 | no | na | na |
| 3L_20774603 | ATTGCAATGTTCACGATGGA | TTGAGTTGCATTGTGGTTCG | 60.3 | 216 | no | na | na |
| 3L_20776277 | CAGGATGGTAGCGATCGAAG | TGCCATCCAATCAGAAAGTG | 60.2 | 228 | yes | TGAGCCTCAATTTGTACNTAACGATGC | TGAGCCTCAATTTGTACNAAACGATGC |
| 3L_20780609 | CCTTTCGGGGAATTTCTGTT | ATGAAACTGCTCCGGATGAC | 60.2 | 223 | no | na | na |
| 3L_20952723 | CGTCAGTGCGGATGAGTAAA | CACTGGTTGGTGAGGTAGCA | 59.8 | 207 | no | na | na |
| 2R_254010 | GAGCATGATCAGAAGCAGCA | AACTGCGTGCAATCCAAGT | 59.8 | 209 | yes | TCTCTGCTCGCTCTCTGCCGCGCATCGTCG | TCTCTACTCGCTCTCTGCCGCGCATCGTCG |
| 2R_273505 | AATGATAACAGCGTGGCAAA | TTCAACCCTACGTGCATCTG | 59.5 | 220 | no | na | na |
| 2R_436659 | ATTGGCCTCGAGCAATGATA | CGGACCATATTTCAGCTCGT | 60.3 | 219 | yes | GCCCCTGCAAGTGGCCAACGCTAACAAACTC | GCCCCTGCAAGTGGCCAACGCTAACAAACCC |
| 2R_494186 | TTCCTTTCAATCAACATGAACG | AGGAGGCTCTCCTAAATGCTG | 60 | 222 | no | na | na |
| 2R_528634 | TTGAAGGCATATTGCATCCTC | TATGCGCTGCACTACGTGAT | 60.3 | 201 | no | na | na |
| 2R_538934 | CATGCCTTTTGTACGGTTGA | TGCCATGGACTAGTGTGCAG | 60.2 | 208 | no | na | na |
| 2R_910700 | TTAGTTTTGGTTCCGCAAGG | CGATCGTCACAGTCGAAAAA | 60 | 220 | no | na | na |
| 2R_5440409 | AAATTTGCTTTTTCGCAAGG | GTCGTTTGACCACATTTTGC | 59.2 | 230 | no | na | na |
| 2R_6752808 | GGCATGACTGTTGTGTTTGG | CGTCGTGGAAAGGTTTTGTT | 60 | 226 | no | na | na |
| 2R_6772242 | TGCTGCACAGACACATACACA | TCTTCTCTTCCTTCCCCACA | 59.9 | 228 | no | na | na |
| 2R_6775346 | ACGGCTTGTCAAAAAGGTTC | GGTTTGTTCGTCCGCATTAT | 59.5 | 230 | no | na | na |
| 2R_6780612 | AAACCAGTTCTGCTGCTGCT | CCATCCGAGAAGGTGAAACT | 59.7 | 227 | no | na | na |
| 2R_6874859 | GGCAATTTTCCCTCCATTTT | TTTGCCCCTAAGCGTGATAC | 60.1 | 211 | no | na | na |

| | | | | | | | |
|---|---|---|---|---|---|---|---|
| 2R_7567517 | TCAATTACCATCGCATGCTC | CACATTTTTATCCCGAAATGG | 59.4 | 229 | no | na | na |
| 2R_7855289 | TGACGAAGATTTTGGTGAAAAA | TAGGCCAGCGAATTTTGTGT | 60.1 | 230 | no | na | na |
| 2R_7856055 | AACGGCCGAGTTTCTGTAAA | GGATCACTGTATTGGGAACGA | 59.8 | 230 | no | na | na |
| 2R_23070067 | CTGTCTTTGTGCATCCCTTG | CGGCACAATTCACCTCACTA | 59.5 | 228 | no | na | na |
| 2R_23431554 | GCACGAGCTTTCTGTGTTCA | TCTACCAGGGCTACCCTTCA | 59.9 | 225 | no | na | na |
| 2R_23437547 | CATCACAATGCGGAGGAAAT | CGCGTACTCTCACCAAACAA | 60.4 | 230 | no | na | na |
| 2R_23463082 | CGCAGCGTAAGAACCACAT | CACAAACCACAACAATGGCTA | 59.7 | 230 | no | na | na |
| 2R_23480856 | CAGCGAGTAGCTGTTGCTCA | GCTTGACCGATGGCTGTAAT | 60.3 | 227 | no | na | na |
| 2R_23481971 | TGCAGATCAATCTTGCCTTG | AACAAAAACGAGCGGATGAG | 60.1 | 217 | no | na | na |
| 2R_23508777 | TTCACCGTCCACCGTAAAAT | CAGCAACACAGAGCTTCGAG | 60.1 | 226 | no | na | na |
| 2R_23534610 | ACGTGTGGAACCTTTGGAAG | GAACCCCGAATGGAAAAGAT | 60.1 | 228 | no | na | na |
| 2R_23539359 | TTGATCAATTGAGCGGTCCT | GCACTTCAACTCCAACAGCA | 60.3 | 218 | no | na | na |
| 2R_23540463 | CCGAGAGACACGAGAGAGGA | TACACCAGCCCACCTACCTC | 60.3 | 220 | no | na | na |
| 2R_23547294 | GAATGCCTTTTCGGATCAAT | TGAAGCAATTGCCAGACATC | 59.4 | 224 | no | na | na |
| 2R_23548651 | CCGGTGTACCTTTCTCCGTA | CCCTTTTCGTCCAAAATGAA | 60 | 221 | no | na | na |
| 2R_23555925 | TGTCTTGCTTCCTGCACACT | GAAAGGGCACCTACGATTCA | 59.8 | 230 | no | na | na |
| 2R_23579923 | AGGATATCGTAACGCGCTTG | GTGGGCCAGAGTAACCAGAG | 60 | 225 | no | na | na |
| 2R_23583342 | CTTGGAATGCGACTCGATCT | GCTTCATACTTTCCGGCTGA | 60.4 | 230 | no | na | na |
| 2R_23590996 | ACCGAGCTTTAGCGTAGCAA | TATGATGGACAAAGCGCATC | 59.9 | 229 | no | na | na |
| 2R_23594562 | CTTACAGCGCCTCCAACATT | CTGATCGGTGTTAGCCACCT | 60.2 | 226 | no | na | na |
| 3R_13040298 | ACCGGTACGTGGTGCTAATC | GTTGTTTGCGGATGCTGTAA | 59.8 | 207 | no | na | na |
| 3R_14783249 | AACCAGCACCAGCAGAAGAG | CCCAGAAATGCAAGTGCTTC | 60.7 | 226 | no | na | na |
| 3R_14848219 | CTCGTGAGAAAGGGACAAGG | TCCCCCTCTGAAAAGGATTC | 60.1 | 169 | no | na | na |
| 3R_14861699 | GGAAACGAATAGGGGAAACC | TCTGCTAACGCACACACACA | 59.9 | 228 | no | na | na |
| 3R_16115639 | CAAAGGGTGATGATGGTGAA | CTGTACCACGAGTTCGGACA | 59.5 | 220 | no | na | na |
| 3R_16134008 | CGACTATGGAGGAACCAGGA | CAAGCTGCAGGGTTCAAGAT | 60.2 | 227 | no | na | na |

Supplementary Table 2. Results of *dadi* analysis for three models.

| **Parameter** | **GOUNDRY admixed (best model)** | **GOUNDRY sister to AT** | **GOUNDRY sister to *A. coluzzii*** |
|---|---|---|---|
| Log likelihood | -83,303.1 | -129,641.9 | -151,999.6 |
| θ | 19,795.7 | 16,498.1 | 11,078.1 |
| Ancestral $N_e$ (calculated from θ) | 818,961 | 682,536 | 458,309 |
| # parameters | 18 | 17 | 17 |
| % of GOUNDRY derived from AT | 83.3% | Constrained to 100% | Constrained to 0% |
| AT $N_e$ early | 1855 | 206,014 | 45,308 |
| AT $N_e$ middle | 1,839,911 | 1,993,450 | 485,599 |
| AT $N_e$ recent | 41,391 | 26,164 | 30,283 |
| *A. coluzzii* $N_e$ early | 4,064,188 | 4,580,735 | 2,971,999 |
| *A. coluzzii* $N_e$ middle | 10,591,123 | 6,156,679 | 9,833,742 |
| *A. coluzzii* $N_e$ recent | 16,836,097 | 33,726,731 | 27,677,151 |
| GOUNDRY $N_e$ | 82,760 | 31,370 | 469 |
| Generations since time of split | 2,238,138 | 1,678,825 | 3,794,845 |
| Generations since first $N_e$ change | 456,499 | 411,755 | 612,717 |
| Generations since second $N_e$ change | 47,989 | 150,795 | 30,338 |
| Generations since GOUNDRY origin | 15,350 | 11,461 | 294 |
| Migration rate *A. coluzzii* to AT, early | 1.7e-08 | 7.9e-07 | 7.4e-07 |
| Migration rate AT to *A. coluzzii*, early | 1.2e-06 | 8.4e-07 | 1.3e-06 |
| Migration rate *A. coluzzii* to AT, middle | 4.3e-06 | 2.4e-04 | 2.5e-06 |
| Migration rate AT to *A. coluzzii*, middle | 1.6e-08 | 2.1e-06 | 4.1e-08 |
| Migration rate *A. coluzzii* to AT, recent | 1.9e-06 | 2.0e-05 | 3.7e-06 |
| Migration rate AT to *A. coluzzii*, recent | 5.9e-07 | 8.5e-08 | 4.9e-06 |
